## Supplemental Figures for "DLG1 functions upstream of SDCCAG3 and IFT20 to control ciliary targeting of polycystin-2"

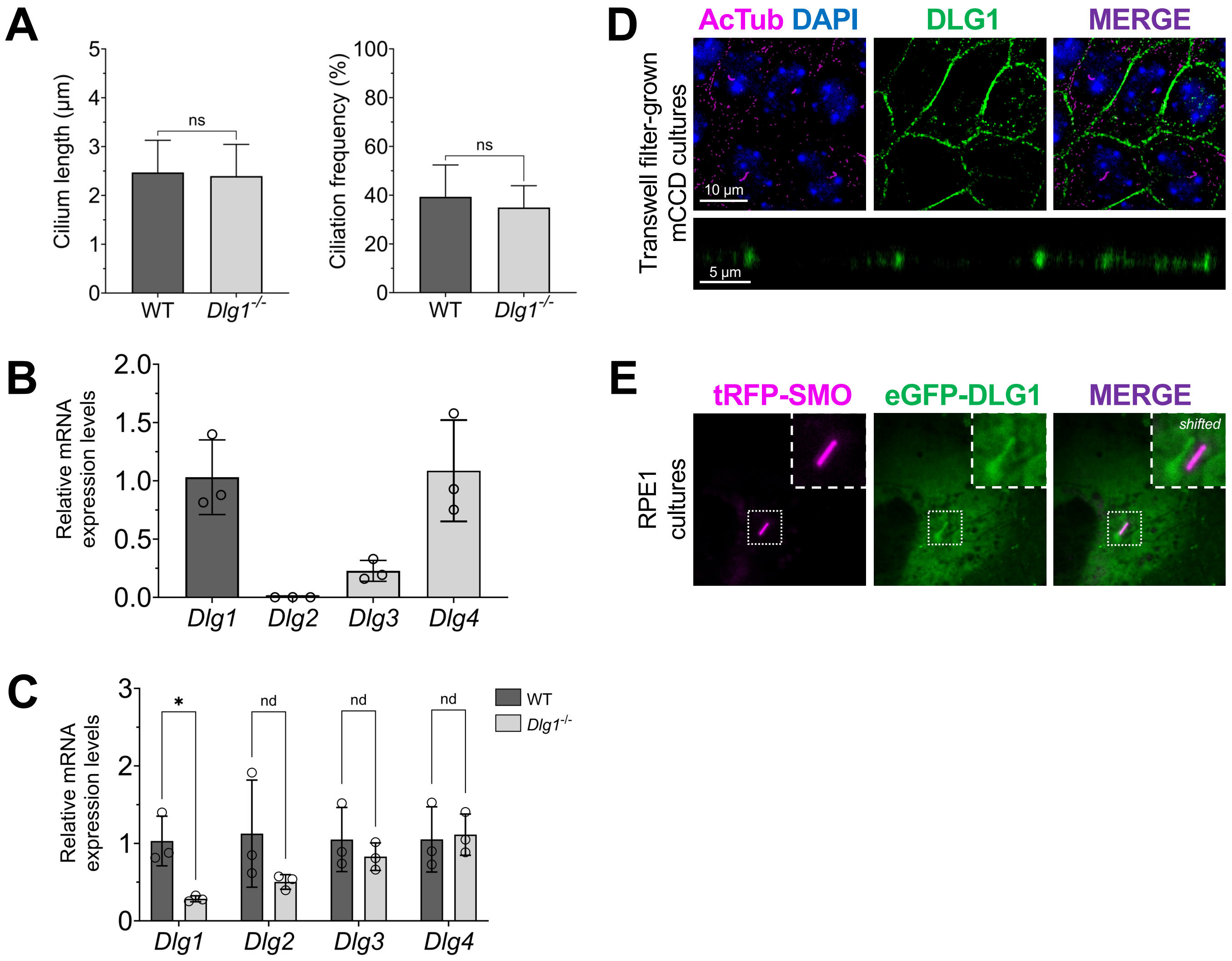


**Figure S1. Characterization of mCCD cell lines and subcellular localization of DLG1.** (**A**) Quantification of ciliary length (left panel) and frequency (right panel) in standard WT and *Dlg1*$^{-/-}$ mCCD cultures. Ciliary length (horizontally oriented cilia) was measured using Fiji. Graphs represent accumulated data (n=3), and statistical analysis was performed using Mann-Whitney U test (unpaired, two-tailed). Error bars represent means ± SD. ns, not statistically significant. (**B**, **C**) Relative mRNA levels of *Dlg1-4* in WT and *Dlg1*$^{-/-}$ mCCD cells were assessed by qRT-PCR. In (**B**) mRNA levels of *Dlg2-4* paralogs in WT cells are normalized to *Dlg1;* in (**C**) *Dlg1-4* mRNA levels in *Dlg1*$^{-/-}$ cells are expressed relative to the respective levels in WT cells. The expression levels are shown as mean with SD error bars of (n=3). For statistical analysis in panel (**C**), multiple unpaired t-test with a two-stage step-up (Benjamini, Krieger, and Yekutieli) method was used. *, $P<0.05$ and nd, not a discovery. (**D**) Localization of endogenous DLG1 (green) in transwell filter-grown mCCD cells. Cilia were stained with acetylated tubulin antibody (magenta) and nuclei stained with DAPI (blue). Upper and lower panels show top and side views, respectively, of reconstructed 3D confocal images of the cells. (**E**) Localization of eGFP-tagged DLG1 (eGFP-DLG1, green) transiently expressed in RPE1 cells stably expressing tRFP-tagged Smoothened (SMO-tRFP, magenta) and visualized by live-cell imaging. Insets show enlarged images of primary cilium and the channels in the merged inset are shifted to aid visualization.

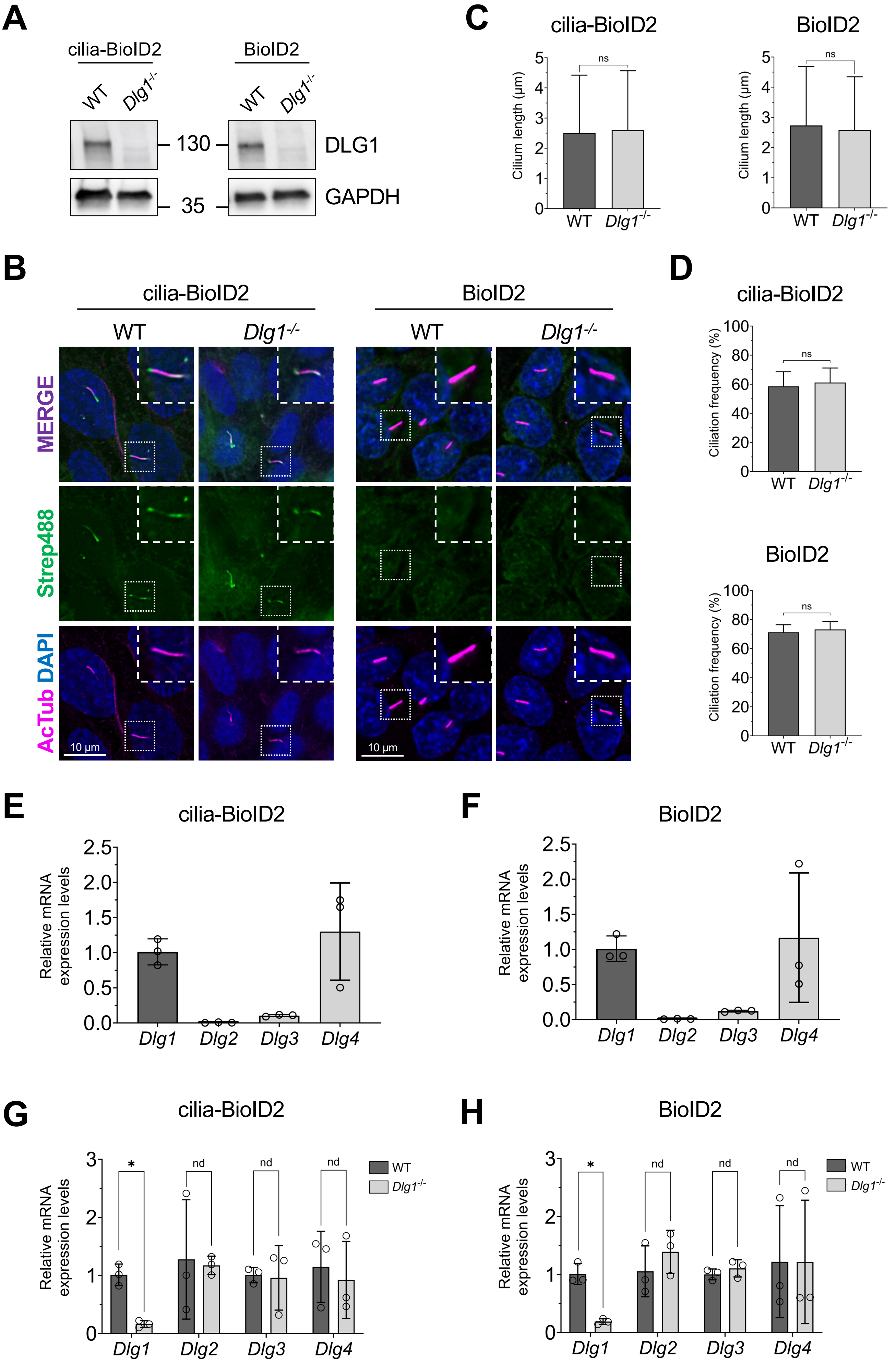

A
cilia-BioID2
BioID2
WT
Dlg1-/-
130
35
DLG1
GAPDH
B
cilia-BioID2
BioID2
WT
Dlg1-/-
MERGE
Strep488
AcTub DAPI
10 µm
C
cilia-BioID2
BioID2
Cilium length (µm)
ns
D
cilia-BioID2
BioID2
Ciliation frequency (%)
ns
E
cilia-BioID2
F
BioID2
Relative mRNA expression levels
Dlg1
Dlg2
Dlg3
Dlg4
G
cilia-BioID2
H
BioID2
*
nd
WT
Dlg1-/-

**Figure S2. Characterization of WT and *Dlg1$^{-/-}$* IMCD3 Flp-In cell lines stably expressing cilia-BioID2 or BioID2.** (**A**) Western blot analysis of indicated cell lines using antibodies against DLG1 and GAPDH (loading control). Molecular mass markers are shown in kDa. (**B**) Representative IFM images of serum-deprived IMCD3 Flp-In WT and *Dlg1$^{-/-}$* cell lines stably expressing cilia-BioID2 (right panels) or BioID2 (left panels), stained with anti-acetylated α-tubulin (AcTub) as cilium marker (magenta), and conjugated streptavidin antibody (Strep488, green) to reveal the biotinylated ciliary proteins; cell nuclei were visualized with DAPI staining. Insets show enlargement of selected cilia. (**C, D**) Quantification of the cilium length (**C**) and the percentage of ciliated cells (**D**) in WT and *Dlg1$^{-/-}$* cells based on images shown in (**B**). Ciliary length (horizontally oriented cilia) was measured using Fiji. Mann-Whitney U test (unpaired, two-tailed) was used for the statistical analysis. Graphs in (**C**) and (**D**) represent accumulated data from three individual experiments. Error bars represent means ± SD. ns, not statistically significant. (**E-H**) Relative mRNA levels of *Dlg1-4* in cilia-BioID2 (**E, G**) and BioID2 (**F, H**) WT and *Dlg1$^{-/-}$* cells were assessed by qRT-PCR. In panels (**C**) and (**F**) *Dlg2-4* paralogs are expressed relative to *Dlg1*, while in panels (**G**) and (**H**) *Dlg1-4* are expressed relative to that in WT cells. The expression levels are shown as mean with SD error bars (n=3). For statistical analysis in panels (**G**) and (**H**), multiple unpaired t-test with a two-stage step-up (Benjamini, Krieger, and Yekutieli) method was used. *, $P<0.05$ and nd, not a discovery.

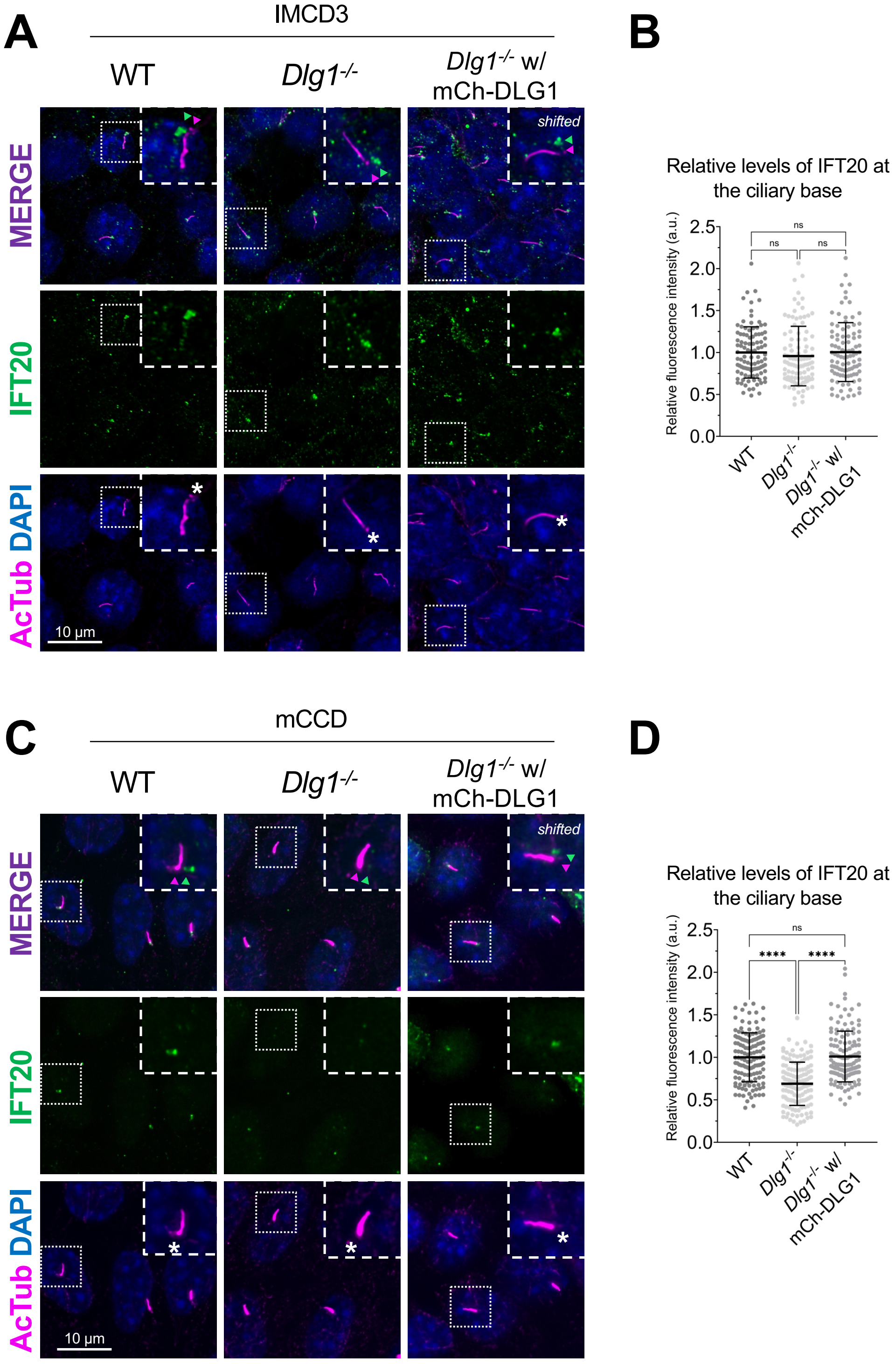


**Figure S3. Ciliary localization of IFT20 in IMCD3 and mCCD cells.** (**A, C**) IFM analysis of ciliated cilia-BioID2 IMCD3 (**A**) and mCCD (**C**) cell lines showing comparative IFT20 staining (green) in WT, *Dlg1*$^{-/-}$ and mCherry-DLG1 (mCh-DLG1) rescue cells. Cilia were stained with antibodies against acetylated α-tubulin (AcTub, magenta), and nuclei visualized with DAPI staining (blue). Insets show enlarged images of cilia, asterisks mark the ciliary base. The merged insets show primary cilia with channels shifted to aid visualization. (**B, D**) Quantification of the relative mean fluorescence intensity of IFT20 at the ciliary base of cilia-BioID2 IMCD3 cell lines (**B**) or at the ciliary base of mCCD cell lines (**D**). Graphs represent WT normalized and accumulated data from three individual experiments. Kruskal-Wallis test with Dunn's multiple comparison test was used for the statistical analysis. Data are shown as mean ± SD. ns, not significant; ****, P<0.0001.

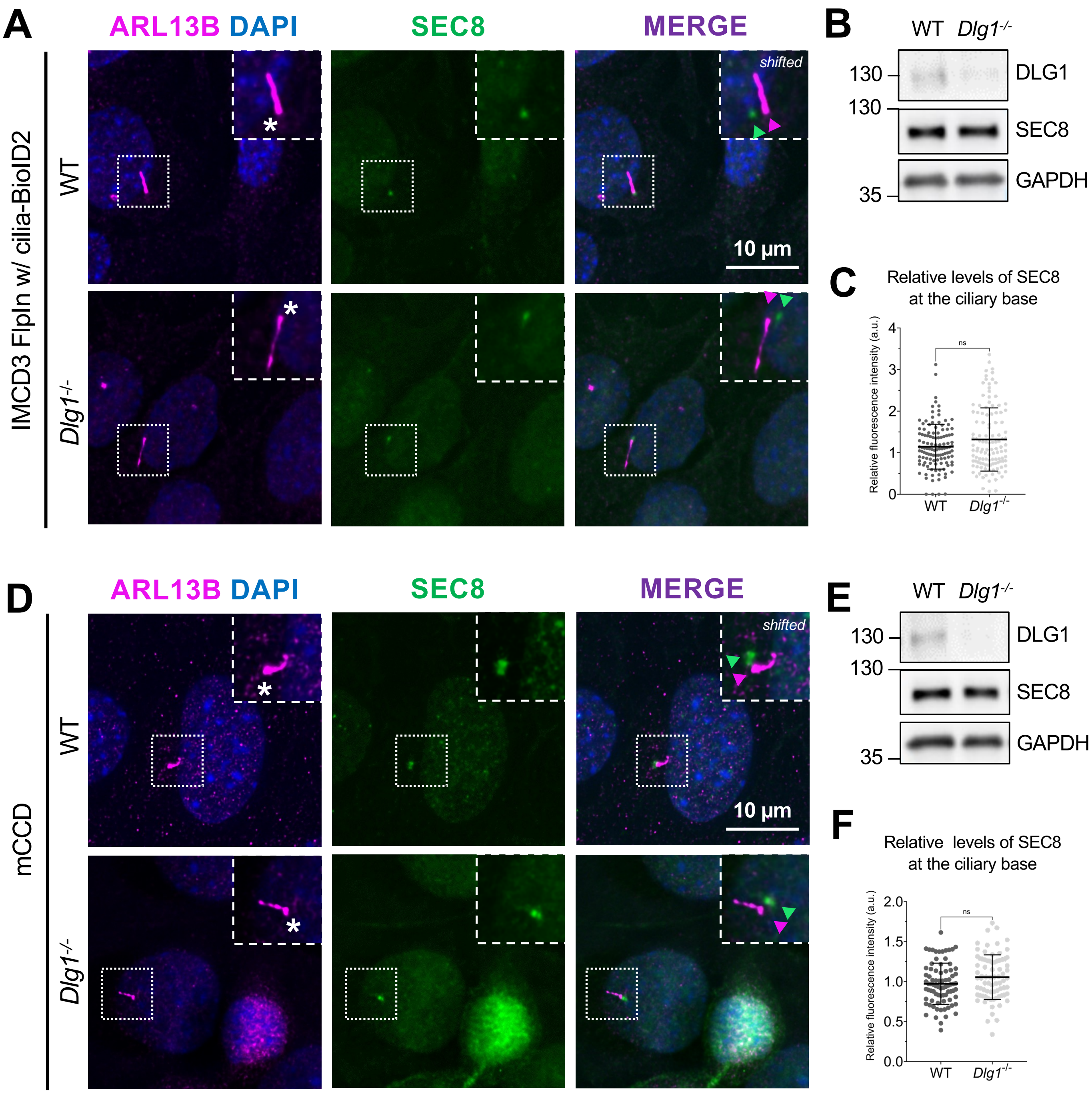


**Figure S4. Loss of DLG1 does not affect ciliary localization of SEC8 in IMCD3 and mCCD cells.** (**A, D**) IFM analysis of ciliated WT and *Dlg1*$^{-/-}$ IMCD3 cilia-BioID2 (**A**) and mCCD (**D**) cells using antibodies against SEC8 (green) and ARL13B (magenta). DAPI was used to mark the nucleus (blue). Insets show enlargement of selected cilia, and the ciliary base is marked with an asterisk. Images of primary cilia in merged channels were shifted to aid visualization. (**B, E**) Western blot analysis of total cellular levels of SEC8 in IMCD3 cilia-BioID2 (**B**) and mCCD (**E**) cell lines. (**C, F**) Quantification of the relative mean fluorescence intensity of endogenously stained SEC8 at the ciliary base in IMCD3 cilia-BioID2 (**C**) or mCCD (**F**) cells. Graphs in (**C**) and (**F**) represent normalized and accumulated data (n=3). Student's t-test (two-tailed, unpaired) was used for the statistical analysis. Data are shown as mean ± SD. ns, not statistically significant.

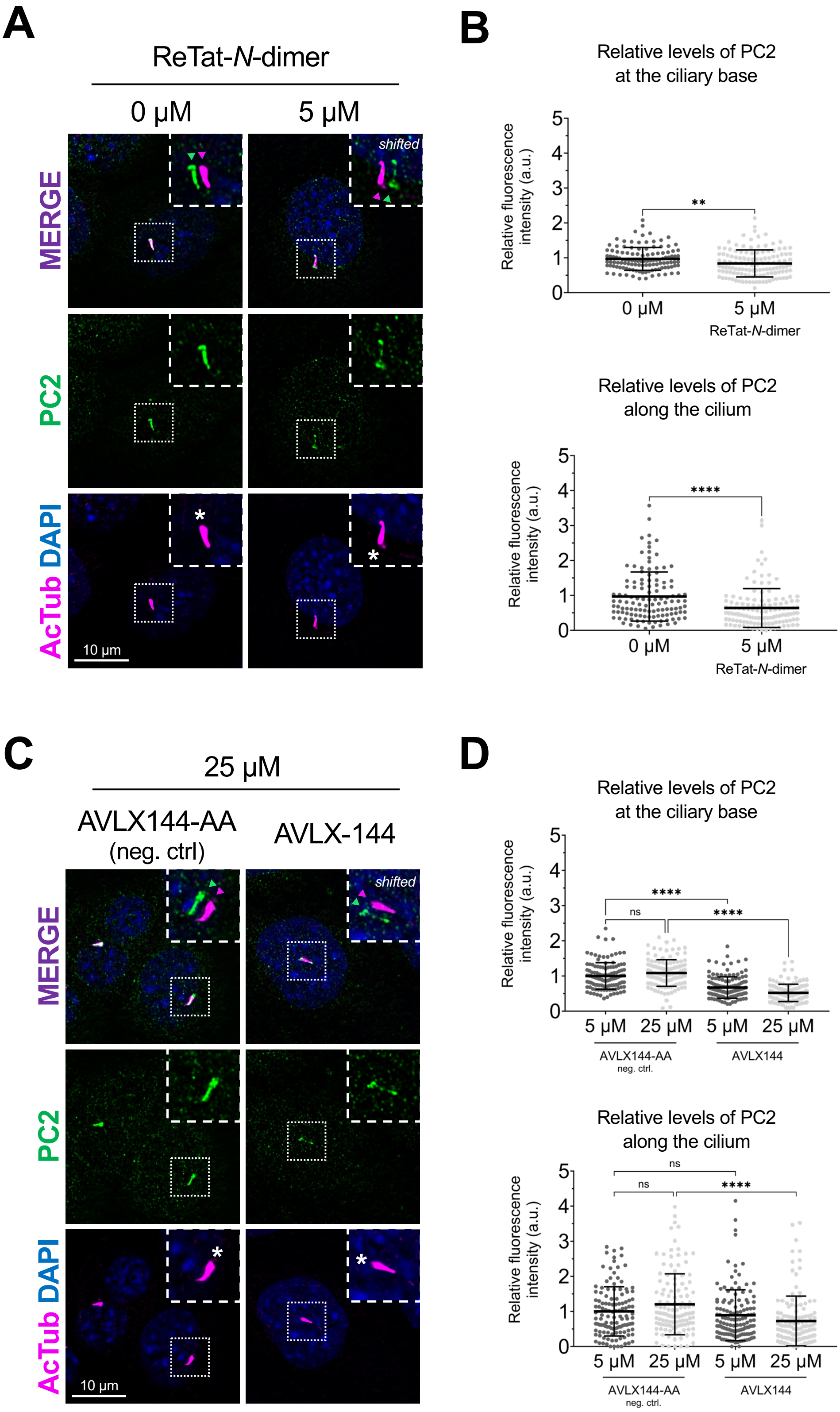
A
ReTat-N-dimer
0 µM
5 µM
shifted
MERGE
PC2
AcTub DAPI
10 µm
B
Relative levels of PC2
at the ciliary base
Relative fluorescence
intensity (a.u.)
**
0 µM
5 µM
ReTat-N-dimer
Relative levels of PC2
along the cilium
****
C
25 µM
AVLX144-AA
(neg. ctrl)
AVLX-144
D
ns
5 µM
25 µM
AVLX144-AA
neg. ctrl.
AVLX144

**Figure S5. Acute DLG inhibition disrupts the localization of PC2 to the primary cilium in mCCD cells.** (**A, C**) PC2 immunofluorescent staining (green) of mCCD cells treated with ReTat-*N*-dimer (**A**) or AVXL144 (Tat-*N*-dimer) (**C**) PDZ inhibitor for 12 h, co-labelled with acetylated α-tubulin antibody (AcTub, magenta) as primary cilium marker and DAPI (blue) as nuclei marker. Insets show enlargements of selected cilia, and the ciliary base is marked with an asterisk. Images of primary cilia in merged channels were shifted to aid visualization. (**B, D**) Quantification of relative PC2 levels at the ciliary base or along the cilium in the indicated conditions. The scatter plot shows the MFI of PC2 at the ciliary base and along the cilium after 12 h of inhibitor treatment with ReTat-*N*-dimer (**B**) or AVLX-144 (Tat-*N*-dimer) (**D**) compared to non-treated cells or treated with the negative control AVXL144-AA, respectively. Graphs represent normalized and accumulated data (n=3). Kruskal-Wallis test with Dunn's multiple comparison test was used for the statistical analysis. Data are shown as mean ± SD. ns, not significant; **, P<0.01; ****, P<0.0001.

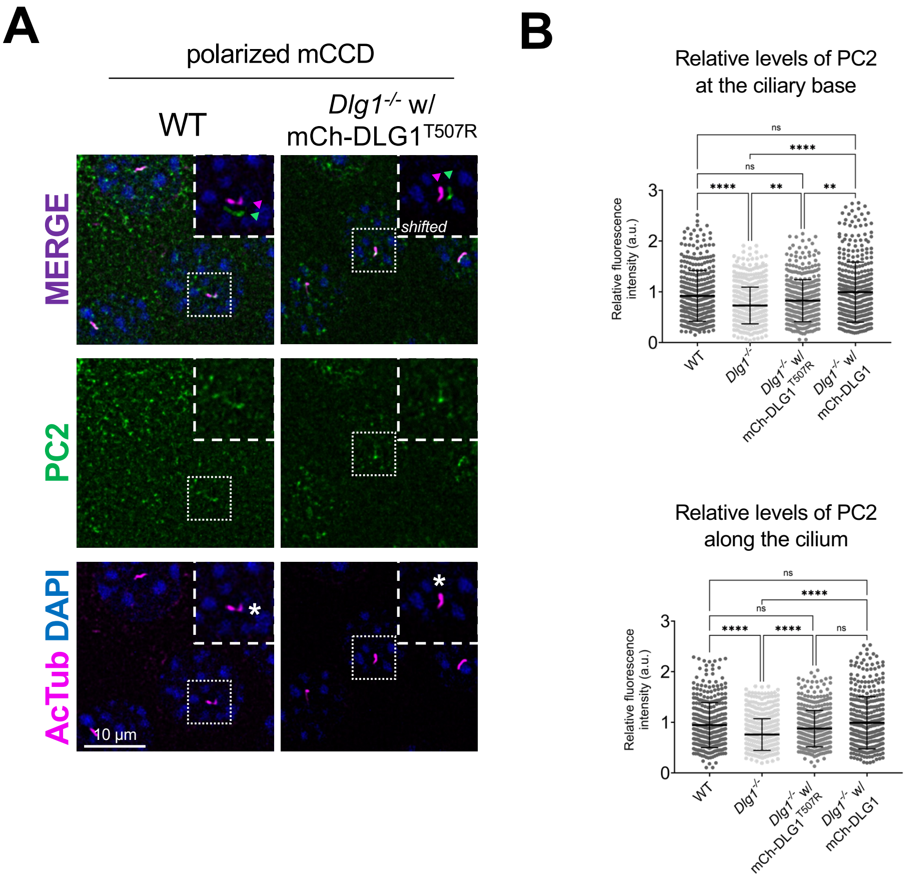


**Figure S6. Effect of a CAKUT-associated DLG1 variant on ciliary PC2 localization mCCD cells.** (**A**) IFM analysis indicated mCCD cell lines cultured on transwell filters and stained with antibodies against PC2 (green) and acetylated tubulin (AcTub; magenta). DAPI was used to mark the nucleus (blue). Insets show enlargement of selected cilia, and the ciliary base is marked with an asterisk. Images of primary cilia in merged channels were shifted to aid visualization. (**B**) Quantification of the relative MFI of endogenously stained PC2 at the ciliary base (top panel) or along the cilium (bottom panel) in the indicated polarized mCCD cell lines. Note that the dataset derives from the same experiments presented in Figure 5C, D; the MFI values for WT, *Dlg1*$^{-/-}$ and *Dlg1*$^{-/-}$ with mCherry-DLG1 (mCh-DLG1) are therefore the same as those shown in Figure 5D. The graphs represent normalized and accumulated data from three individual experiments. Kruskal-Wallis test with Dunn's multiple comparison test was used for the statistical analysis. Data are shown as mean ± SD. ns, not significant; **, P<0.01; ****, P<0.0001.

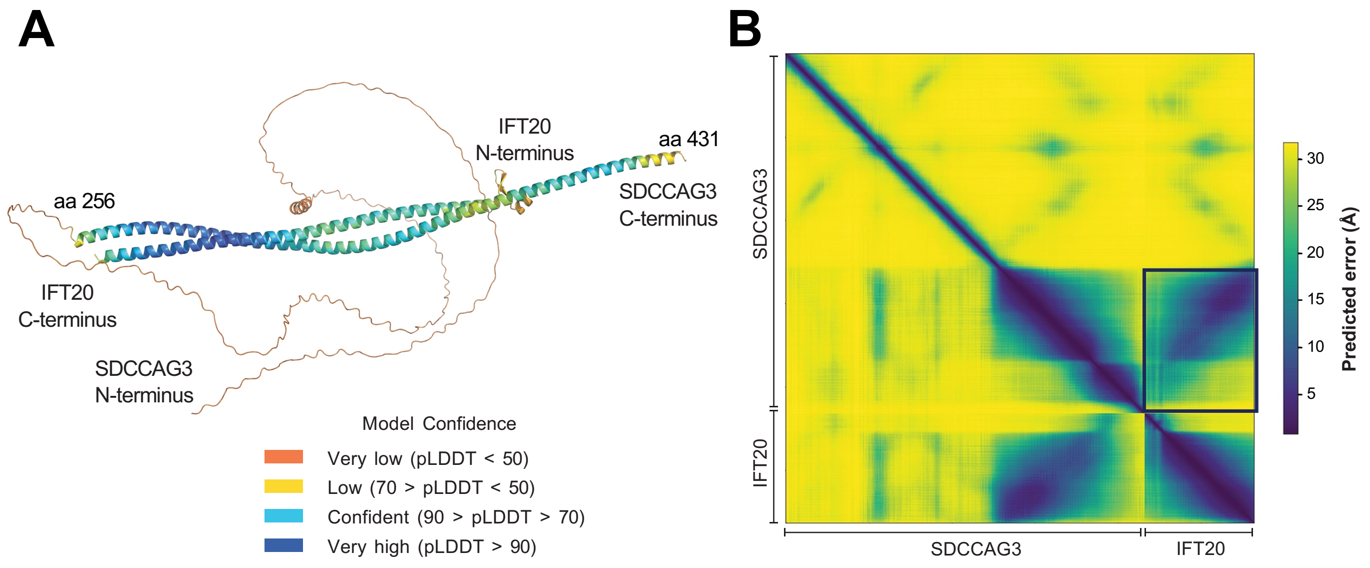


**Figure S7. AlphaFold modelling of SDCCAG3 and IFT20 interaction.** (**A**) AlphaFold prediction of the MmSDCCAG3-IFT20 complex. Amino acids in the complex are color-coded according to their per-residue confidence score (pLDDT), from 0 to 100. Low scores indicate low confidence, and high scores indicate high confidence predictions. (**B**) Predicted alignment error (PAE) plot of MmSDCCAG3-IFT20. The PAE scores for the IFT20-SDCCAG3 interaction are shown within the black box. The boxed area shows low PAE along a diagonal consistent with a high-confidence prediction of an antiparallel heterodimeric coiled-coil complex.

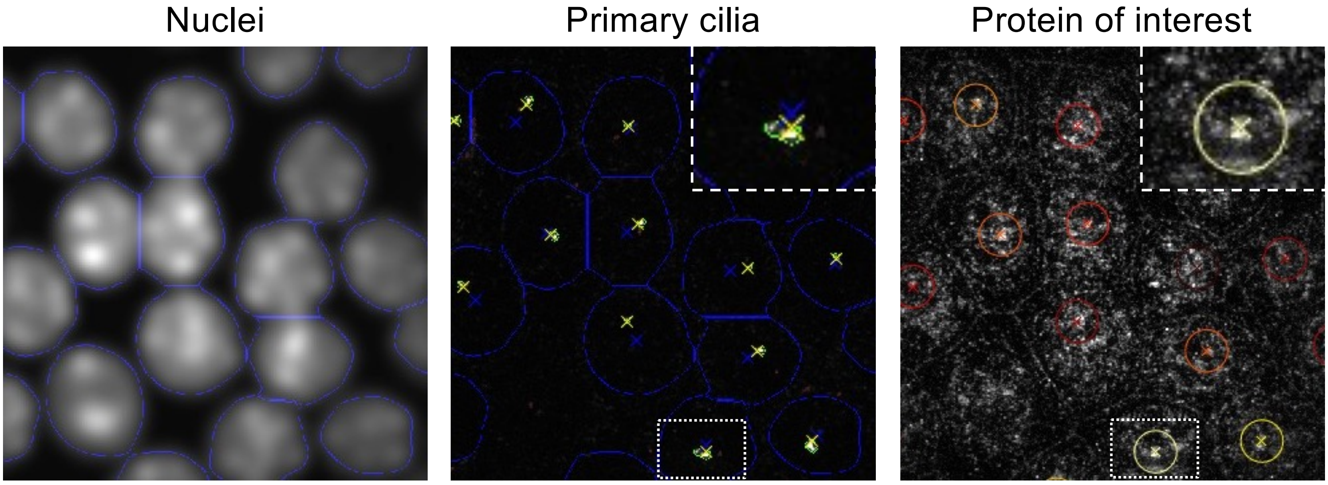


**Figure S8. Overview of the outcome of the MATLAB script-based analysis.** Nuclei segmentation (left) and primary cilia segmentation (middle) defined as the brightest 3D objects overlapping a nucleus region. Primary cilium bases (yellow cross) are identified as the closest cilium voxel to the centroid of the corresponding nucleus region.
