## Supplementary material for "DLG1 functions upstream of SDCCAG3 and IFT20 to control ciliary targeting of polycystin-2": Uncropped western blots

Figure 1D

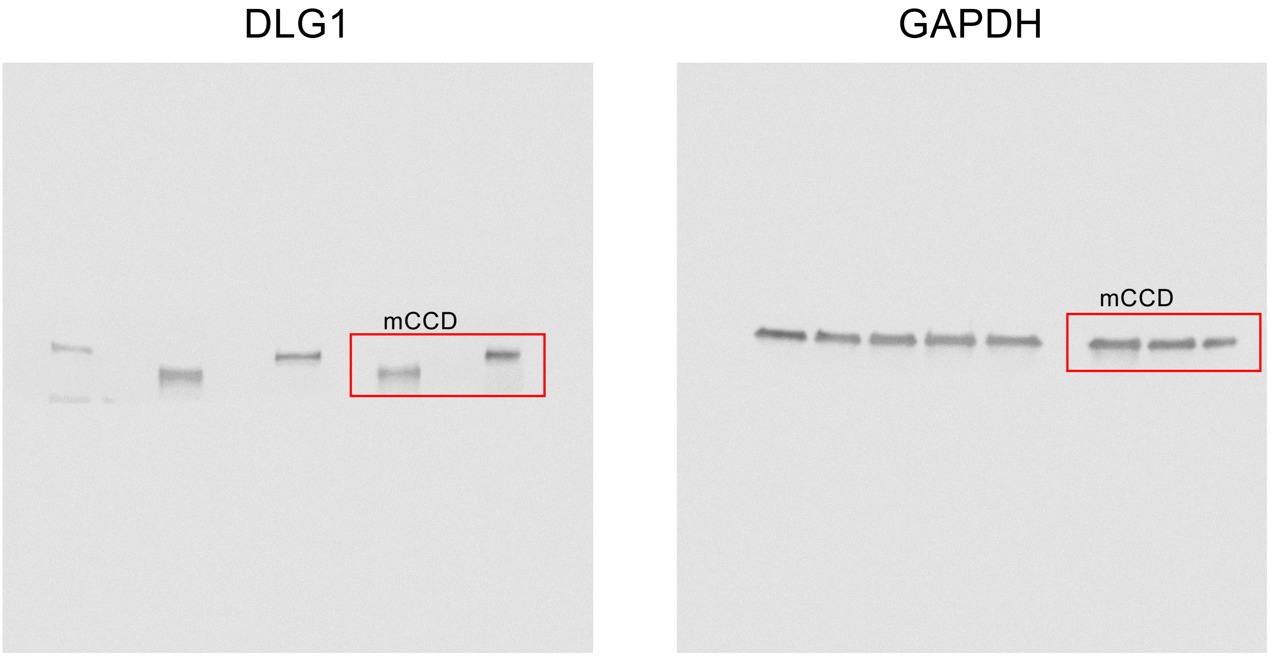

Figure 3C

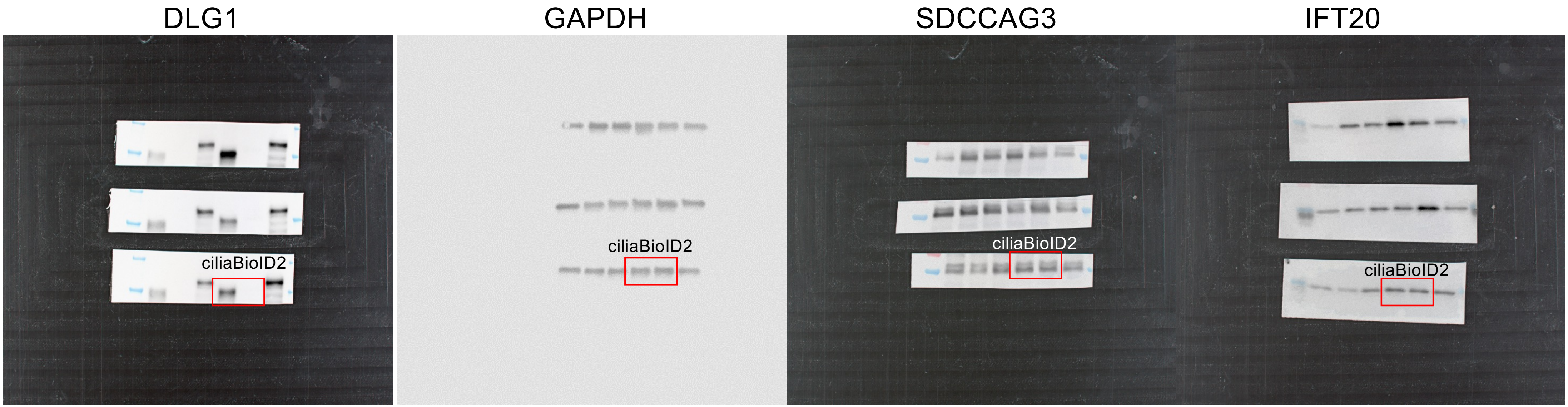

DLG1
GAPDH
SDCCAG3
IFT20
ciliaBioID2
ciliaBioID2
ciliaBioID2
ciliaBioID2

Figure 3D

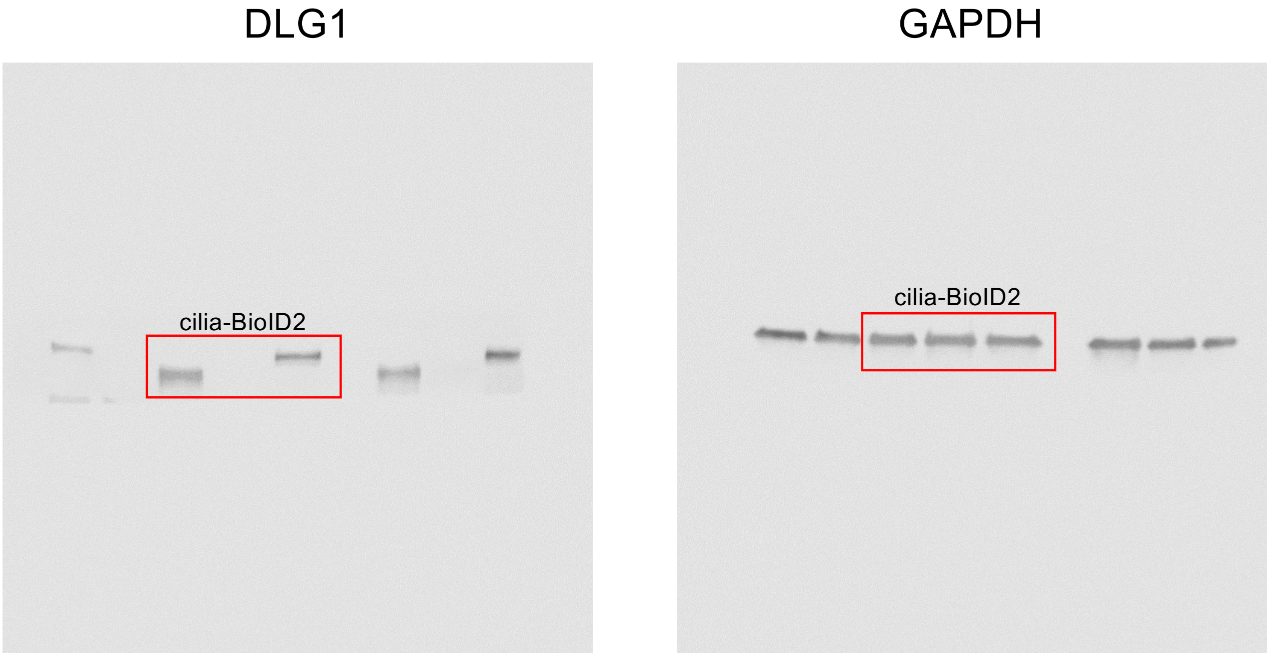

Figure 3G

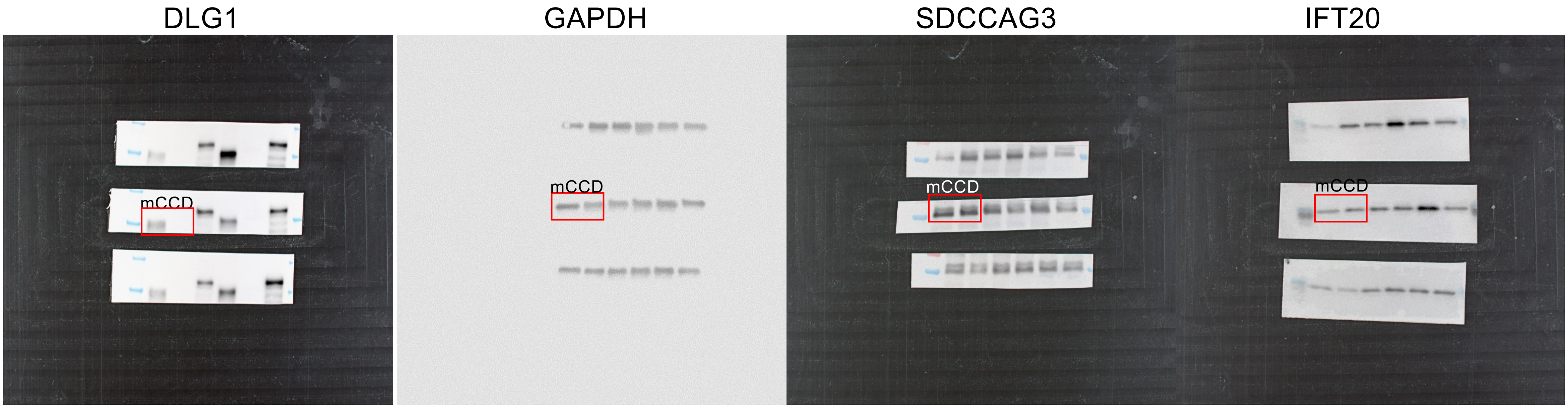

Figure 6B

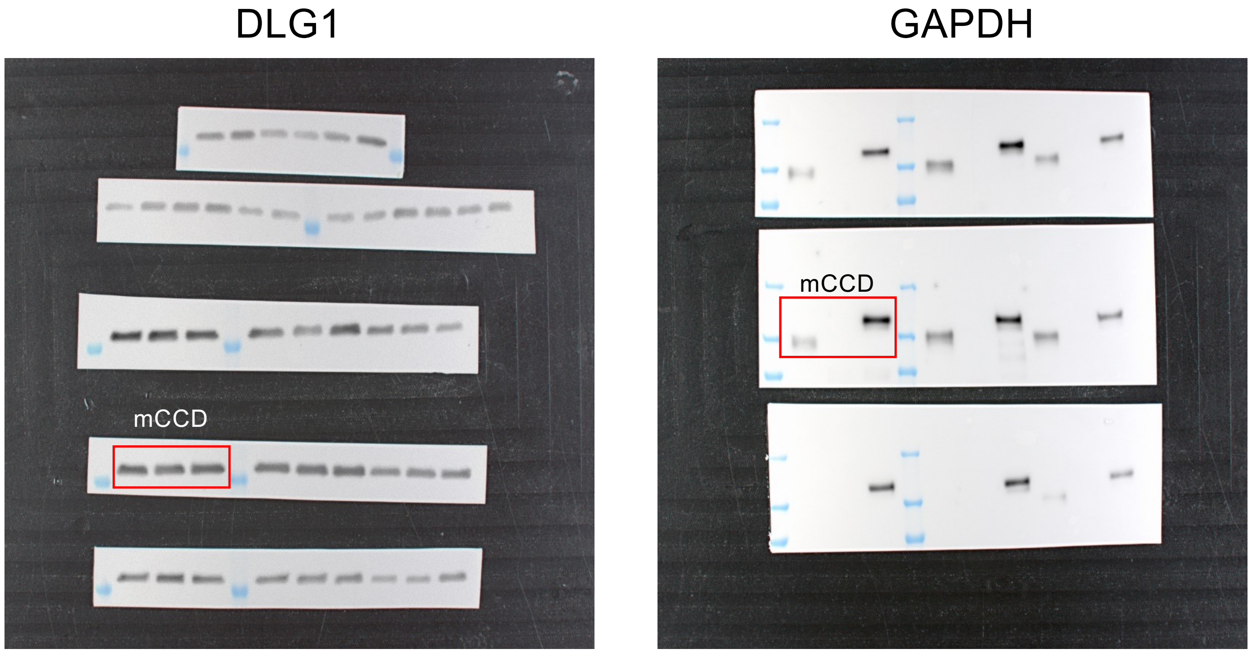

Figure 6G

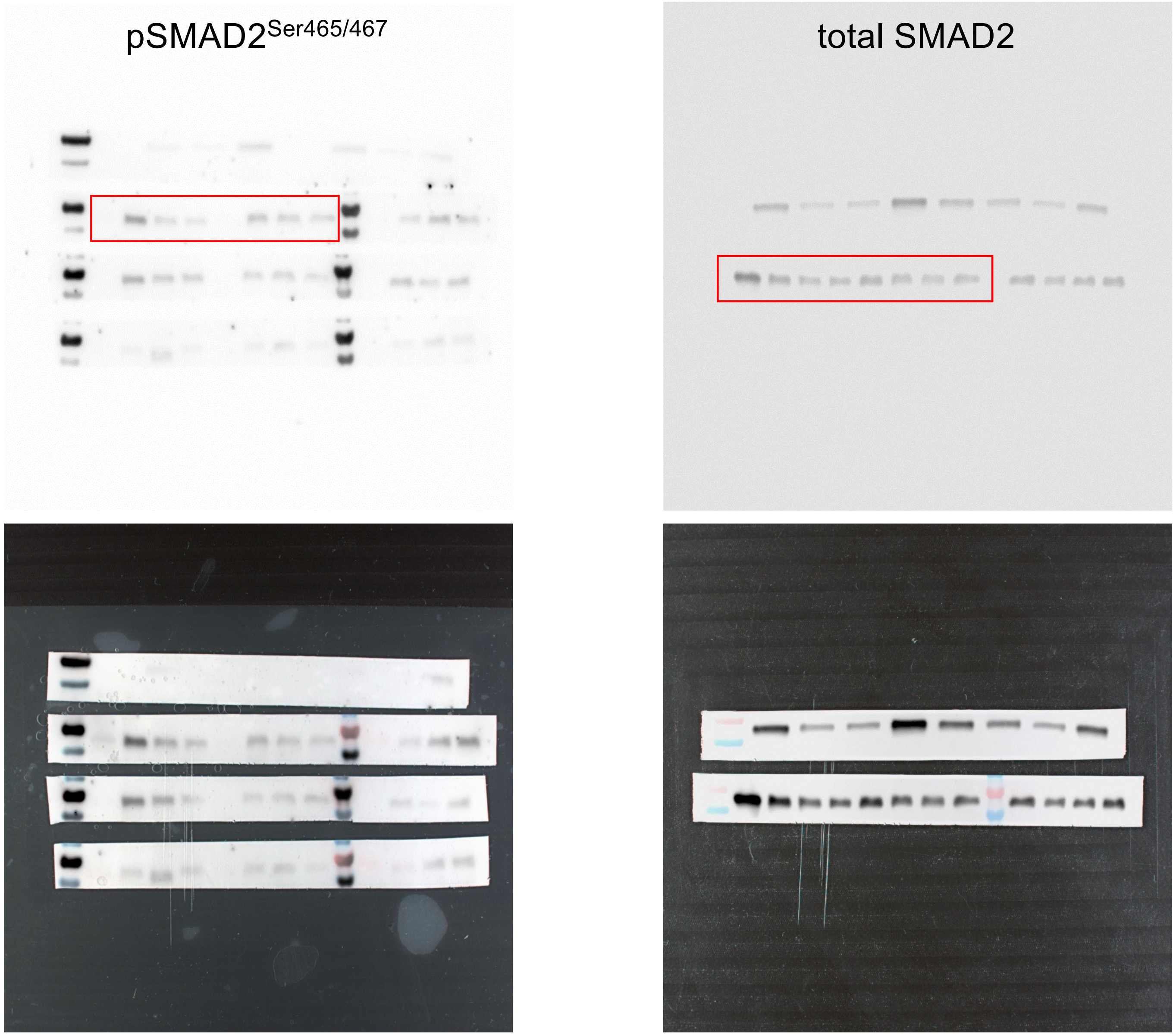

Figure 6I

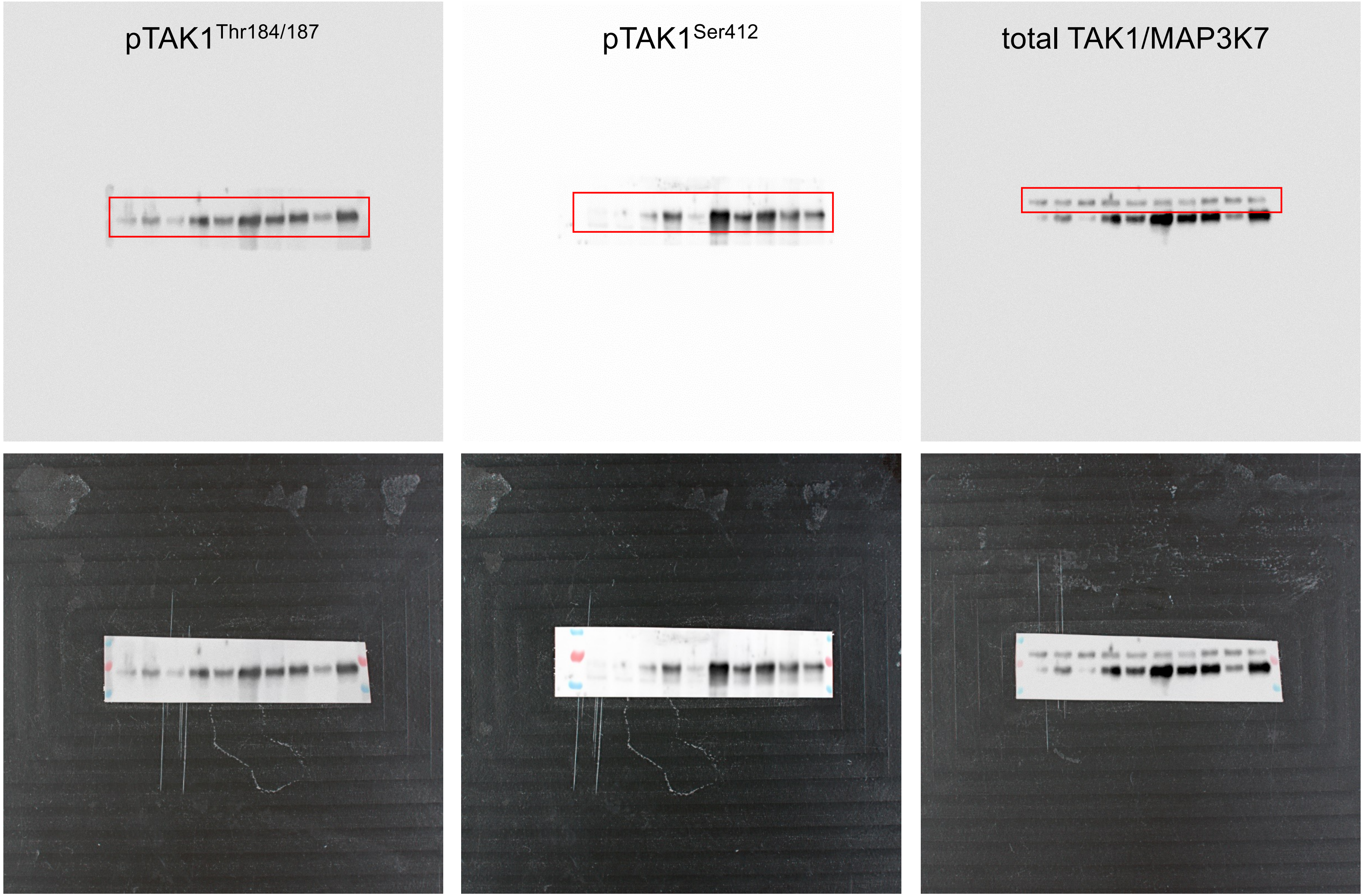

Figure 7A

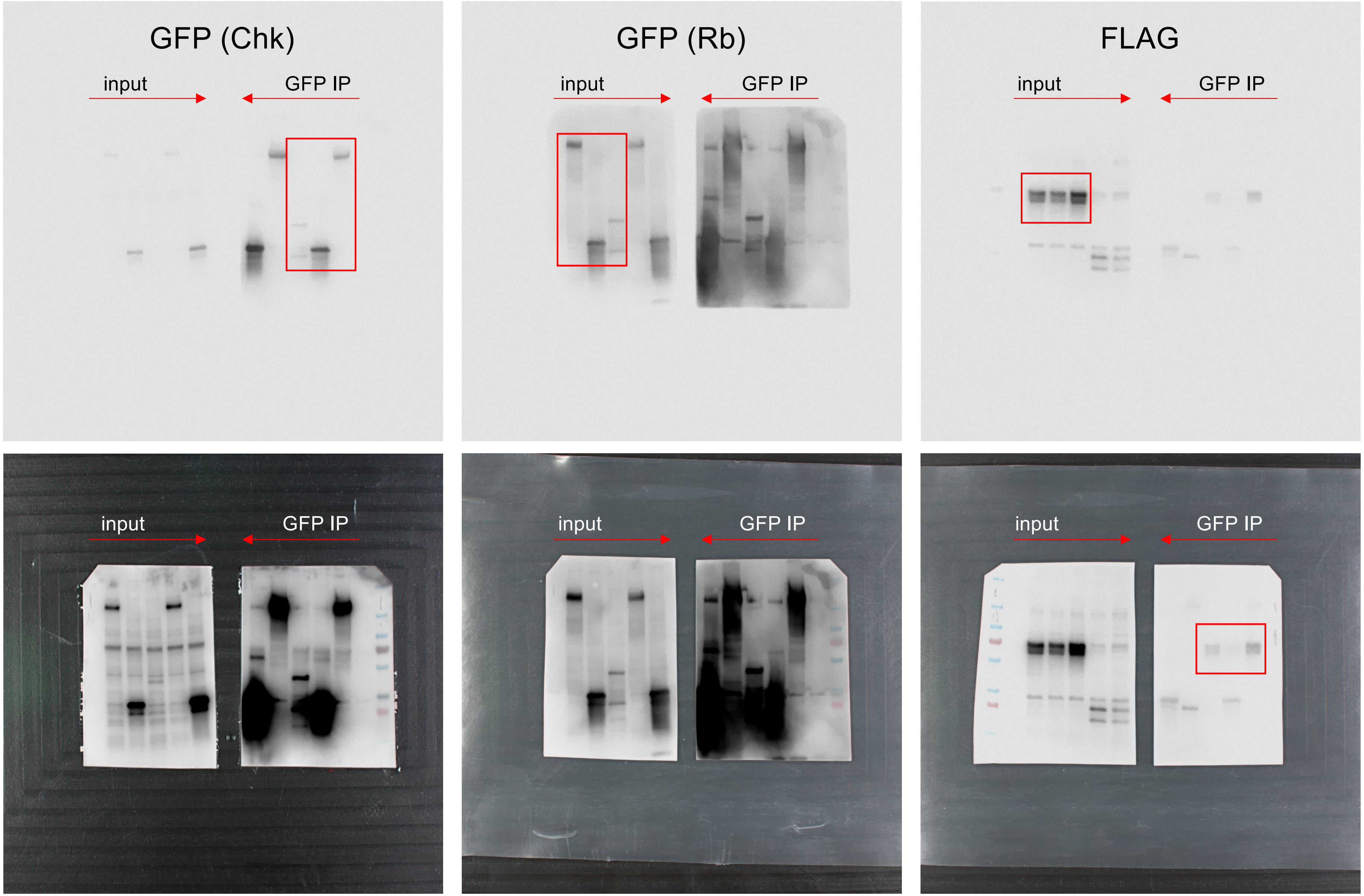
GFP (Chk)
input
GFP IP
GFP (Rb)
input
GFP IP
FLAG
input
GFP IP
input
GFP IP
input
GFP IP
input
GFP IP

Figure 7B

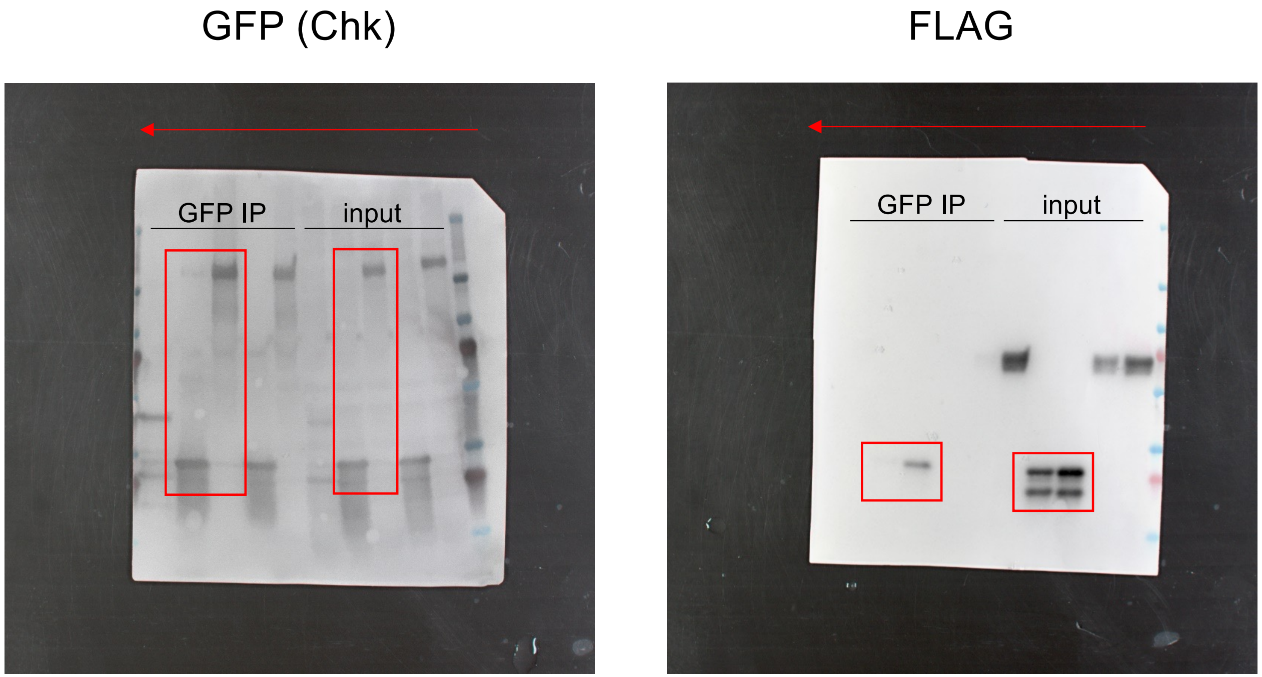

Supplemental Figure 2A

DLG1

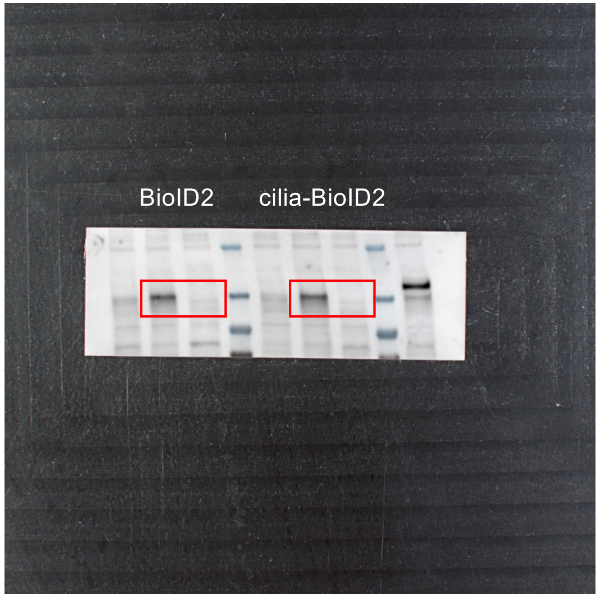


GAPDH

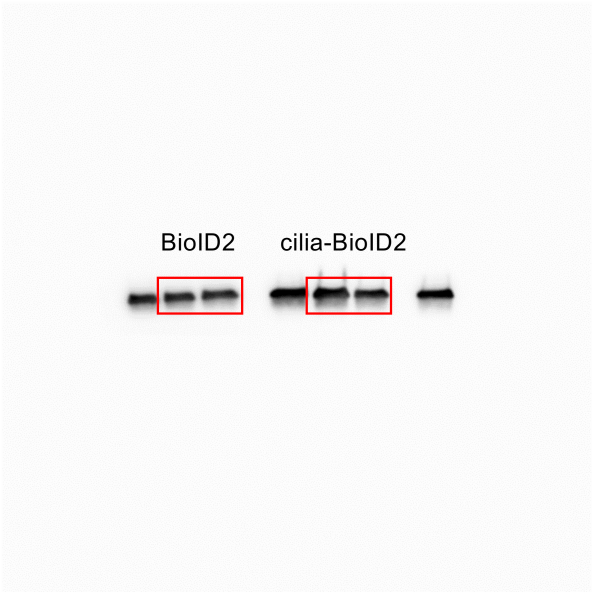

Supplemental Figure 4B and 4E

DLG1

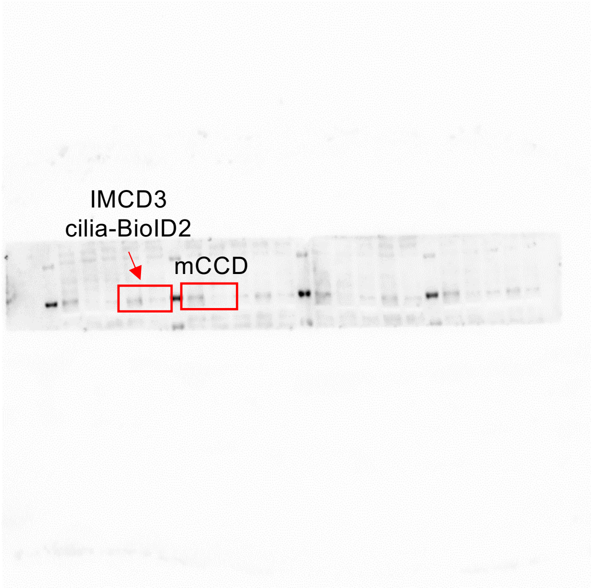


SEC8

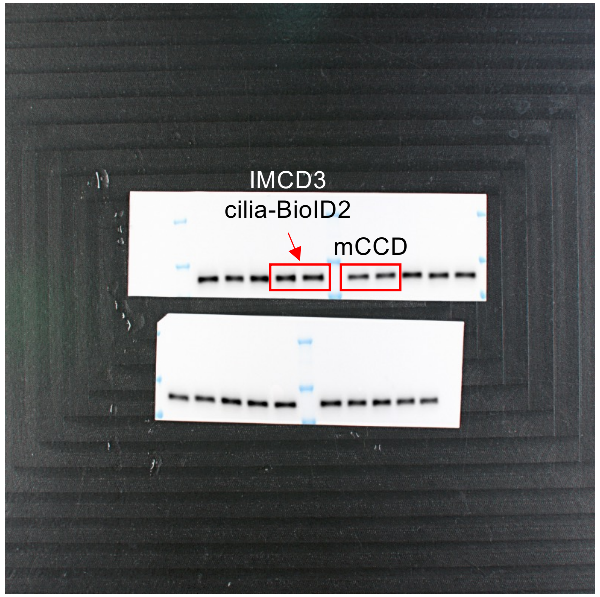


GAPDH

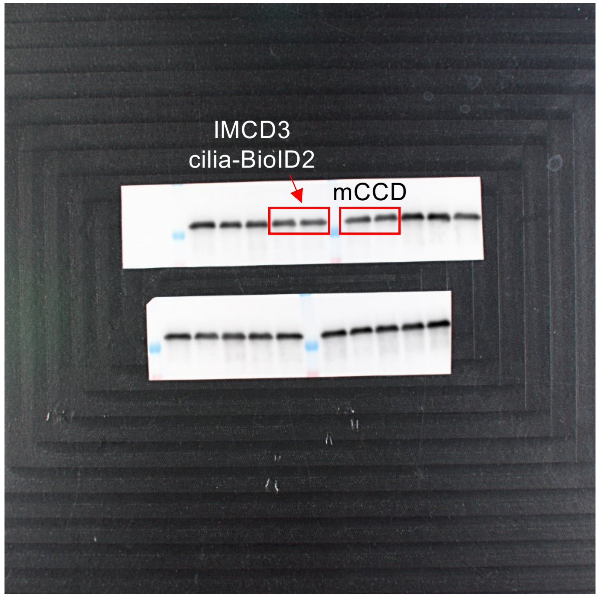
